# Elucidating conformational alteration of human islet amyloid polypeptide by nonsynonymous substitution

**DOI:** 10.1101/2024.11.16.623924

**Authors:** Md Abul Bashar, Largess Barua, Supti Paul, Nayan Dash, Toma Sadhu, Sarmistha Mitra, Raju Dash

## Abstract

Islet amyloid polypeptide (IAPP) is a peptide hormone that serves multiple essential functions, including metabolism and regulating gastric emptying and satiation through amylin receptors. However, mutations in the *IAPP*, notably in the amyloidogenic segment (20-29 amino acid residues), cause its aggregation and amyloid formation, which leads to β-cell toxicity and death in type 2 diabetes mellitus (T2DM) and protein misfolding disorders (PMDs). The current work aims to elucidate the non-synonymous variants in the *IAPP*, which may adversely affect its function and rise to T2DM and PMDs. We harnessed *in silico* non-synonymous single-nucleotide polymorphisms (nsSNPs) assessment and molecular dynamics (MD) simulation to discover the potential deleterious mutants that cause T2DM and PMDs. Firstly, we executed nsSNPs prediction in *IAPP* using the NCBI dbSNP server, and then, all the predicted nsSNPs were assessed by a total of 26 *in silico* tools to find out which possessed the most deleterious effect in IAPP. Finally, MD simulation was carried out utilizing the most deleterious nsSNPs to check which significantly alters the conformational dynamics of IAPP. We found a total of 62 nsSNPs, among which the top 4 deleterious nsSNPs (T37P, L45P, G66R, and T69I) were selected based on the deleteriousness predictions by in silico tools and their location in the mature *IAPP* sequence (34-70 amino acid residues). MD simulations further confirm that three variants (T37P, L45P, and G66R) significantly alter the conformational dynamics of IAPP, suggesting a potential starting point for future research to elucidate the roles of these variants in IAPP aggregation and amyloid formation and their associations with T2DM and PMDs.

## 1 Background

Islet amyloid polypeptide (IAPP), also referred to as amylin, is a peptide hormone consisting of 37 amino acids that is produced in the pancreas in response to various foods such as lipids, glucose, or amino acids (such as arginine) [1, 2]. It exerts an essential role in metabolism, notably glucose homeostasis, by reducing the secretion of nutrient-stimulated glucagon and regulating gastric emptying and satiation [3-7]. It also inhibits osteoclast activity, enhances osteoblast proliferation [8, 9], and regulates blood pressure by affecting the renin-angiotensin system [10]. The gene responsible for IAPP expresses a pre-pro-hormone (called pre-pro-]IAPP) consisting of 89-amino acid that is later on converted to pro-IAPP consisting of 67-amino acids (residues, 23-89) by signal peptidase that removes signal peptide (1-22 residues) and a disulfide bond is formed between two Cys residues of at 2 and 7 positions, catalyzed by protein disulfide isomerase (PDI) [11, 12]. Prohormone convertase 2 (PC2) and prohormone convertase 1/3 (PC1/3) removes 11 amino acids from N-terminus and 16 amino acids from C-terminus of pro-IAPP [13-15], followed by the removal of C-terminal dipeptide Lys–Arg (residues, 72-73) by carboxypeptidase E (CPE) and subsequently peptidylglycine α-amidating monooxygenase (PAM) removes Gly (residues, 71) and amidates Tyr (residues, 70), leading to the formation of mature IAPP (residues, 34-70) [11, 16-18]. The pancreatic β-cells store and secrete mature IAPP along with insulin[19-21]. IAPP exerts its physiological effects through its receptors (called IAPP or Amylin receptors) that are formed when the calcitonin (CT) receptor is coexpressed with receptor activity-modifying proteins (RAMPs) [22-25]. Nevertheless, its precise mechanism of action through its receptors is not yet fully understood. IAPP also interacts with hormones like insulin [26], glucagon-like peptide-1 (GLP-1) [27], cholecystokinin (CCK) [27, 28], and leptin [29] and modulates blood glucose levels.

Recently, IAPP has been reported to be aggregated and form amyloid, a process called amyloidogenesis, possibly due to the mutations in the *IAPP* sequences that significantly alter the aggregation propensity of IAPP [30, 31]. Previous studies have shown that the amino acids 20–29 residues comprise the central amyloidogenic segment of IAPP [32-34]. For example, a naturally occurring polymorphic S20G mutation renders the human IAPP more aggregation-prone, found with minor frequency in Chinese and Japanese populations, and this mutation is associated with an increased risk of early onset of type 2 diabetes [31]. Additionally, two other regions of IAPP, the carboxyl-terminal region 30–37 [34] and amino acids 8–20 [35], have been identified as potential amyloidogenic regions. For instance, an *in silico* study demonstrated that an F15L variation impairs the α-helix and β-sheet propensities of the protein, resulting in rapid amyloid formation [36]. It has been reported that the deposits of IAPP are found in more than 90% of patients with type 2 diabetes mellitus [37-41], hence a histopathological hallmark of the pancreatic islets, where it is thought to contribute to beta cell dysfunction and death [42-44]. The mechanism by which IAPP causes cytotoxicity and β-cell death (apoptosis) is possibly through membrane permeabilization [45, 46], plasma membrane blebbing, chromatin condensation, and DNA fragmentation [47], increased ER and oxidative stress [48-50], proinflammatory cytokine release and activation [51, 52], increased ROS production [53, 54], mitochondrial dysfunction [55, 56], and advanced glycation end (AGE) products [57]. Recently, IAPP has also been reported to be involved in autophagy dysregulation [58, 59], UPS defects [60, 61], Alzheimer’s [62], and heart disease [63].

Previous studies have confirmed that aggregation of IAPP and amyloid formation is the primary cause of β-cell toxicity and death in T2DM [42-44]. IAPP, when aggregated, can promote the misfolding and formation of soluble oligomers and insoluble amyloid fibrils at increased concentration due to its highly amyloidogenic, intrinsically disordered, and aggregation-prone properties, causing β-cell toxicity and death [42-44, 64]. Additionally, aggregation of misfolded proteins is the primary characteristic of a group of diseases called protein misfolding disorders (PMDs), including several neurodegenerative diseases, such as Alzheimer’s disease (AD), Parkinson’s disease (PD), amyotrophic lateral sclerosis (ALS), and prion disorders, in addition to different systemic amyloidosis diseases [65, 66]. Misfolded protein aggregates are thought to accumulate in the brain and hence cause neurodegenerative disorders [65, 66]. However, the aggregation, misfolding, and amyloid formation, which spread to different body parts and cause toxicities by IAPP, is still poorly understood. Therefore, the elucidation of the mechanism by which IAPP aggregates and forms amyloid may assist in apprehending the pathophysiology of the IAPP-induced β-cells toxicities and PMDs and may be necessary for the development of novel therapeutics (inhibitors) to prevent aggregation of IAPP and hence prevent β-cell toxicity and death in IAPP-mediated type II diabetes.

In this study, we performed *in silico* nsSNPs prediction and MD simulation to discover the most deleterious *IAPP* variants. At first, we found 62 nsSNPs of *IAPP* by NCBI dbSNP web server. These nsSNPs underwent deleteriousness assessment using 26 in silico tools, and based on the results, 4 of the most deleterious nsSNPs were selected. MD simulation further validates 3 variants among those 4 variants (T37P, L45P, and G66R) significantly alter the conformational dynamics of IAPP.

## 2 MATERIALS AND METHODS

### 2.1 Data extraction and prediction of deleterious SNPs

The nonsynonymous single nucleotide polymorphisms (nsSNPs) associated with the *IAPP* gene were extracted from the NCBI dbSNP database along with their corresponding rsIDs [67]. A total of 62 missense SNPs were retrieved, which were subsequently screened utilizing 26 widely recognized bioinformatics tools to predict the most potentially deleterious nsSNPs **(Supplementary File 1)**, including SIFT, FATHMM, M-CAP, MetaLR, MutPred, MutationTaster, PROVEAN, PolyPhen-2 HumDiv and HumVar, VEST4, fathmm-MKL, Condel, PhD-SNP, SNPs&GO, I-Mutant 3.0, CADD-PHRED and CADD-Raw, ClinPred, DANN, MVP, MetaRNN, REVEL, PredictSNP, LIST-S2, PANTHER, PrimateAI, MPC and GenoCanyon. The cut-off of these tools is listed in **Table S1 (Supplementary File 2)**.

SIFT is an *in silico* tool based on the PSI-BLAST algorithm that harnesses sequence homology to estimate the possible impact of amino acid substitutions on protein function, where the predictive value spans from 0 to 1 [68]. Functional analysis through hidden Markov models (FATHMM) is an *in silico* tool designed to predict the functional consequences of coding and noncoding variations, and it can differentiate neutral variations from harmful ones [69]. The clinical pathogenicity classifier M-CAP utilizes a gradient-boosting tree classification model, incorporates 318 characteristics as input, and can detect missense mutations with 95% sensitivity [70]. MetaLR employs logistic regression to incorporate nine distinct deleteriousness values for variants [71]. MutPred is a web-based application that employs a random forest method to assess alterations in protein structures and dynamics resulting from variations while also providing predictions regarding protein functionality, amino acid sequences, and evolutionary data [72]. MutationTaster utilizes a Bayesian classifier to eventually assess the pathogenic potential of a mutation, where the Bayes classifier estimates the functional implications of amino acid changes, intronic and synonymous modifications, short insertion and/or deletion mutations, and variations that traverse intron-exon boundaries [73]. The score ranges from 0 to 1, with higher values indicating a greater likelihood of being harmful [73]. PROVEAN is a predictive technique utilizing sequence clustering to estimate changes in protein function resulting from amino acid substitutions [74]. PolyPhen-2 assesses the functional impact of an allele substitution based on its individual characteristics using a Naïve Bayes classifier [75]. It provides two prediction models: HumDiv and HumVar, where HumDiv primarily found the less deleterious SNPs, but HumVar categorized the SNPs with significant phenotypic impact based on PSIC score [75]. The Variant Effect Scoring Tool 4 (VEST4) is a machine learning classifier utilizing random forest algorithms to identify potentially functional missense variants [76]. fathmm-MKL is a web-based application that integrates functional annotations from ENCODE with nucleotide-level sequence conservation metrics to assess the effects of coding and noncoding variations [77]. Condel is an integrated approach utilizing five predictive techniques to detect missense single nucleotide variations [78]. The integration of five distinct techniques may enhance the prediction of missense mutations’ effects on protein biological activity [78]. PhD-SNP is a technique utilizing support vector machines to estimate if a point mutation is a neutral polymorphism or linked to genetic diseases in people [79]. SNPs and GO provide a framework for determining if a mutation is linked to a disease based on protein sequence analysis [80]. I mutant 3.0 is a computational method utilizing a support vector machine to predict changes in protein stability resulting from single point mutations [81]. Combined Annotation-Dependent Depletion (CADD) is a prevalent machine learning model that integrates over 60 genomic features to assess human single nucleotide variations and brief insertions and deletions throughout the reference assembly [82]. CADD offers two score types: CADD-Raw and CADD-PHRED [82]. ClinPred is a promising tool for detecting disease-associated nonsynonymous variations by integrating two machine-learning models and provides superior accuracy in pathogenicity prediction [83]. DANN assesses every potential single nucleotide variation to elucidate non-linear correlations among features by employing an artificial deep neural network [84]. The score ranges from 0 to 1, with higher values indicating a greater likelihood of being deleterious [84]. MVP (missense variants prediction) is an efficient deep learning approach for predicting the pathogenicity of missense mutations, constructed on a deep residual network (ResNet) model of two blocks [85]. MetaRNN is a deep learningbased technique for predicting pathogenicity that identifies and prioritizes uncommon nonsynonymous single nucleotide variations (nsSNVs) and non-frameshift (NF) indels [86]. REVEL is a prevalent meta-predictor that employs the scores from 13 distinct *in silico* tools to assess the pathogenicity of missense mutations [87]. The REVEL score for missense variants ranges from 0 to 1, with elevated values indicating an increased probability of the variant being pathogenic [87]. PredictSNP tool is a consensus SNP classifier that utilizes six different *in silico* tools to predict disease-associated mutations [88]. LIST-S2 is a web-based tool that utilizes local sequence identity and taxonomic distances to estimate the deleteriousness of mutations [89]. Protein ANalysis THrough Evolutionary Relationships (PANTHER) is an *in silico* tool that produces a substitution position-specific evolutionary conservation score (subPSEC) by employing multiple sequence alignments and the Hidden Markov Model (HMM) [90]. SNPs with a value below -3 are deemed harmful, but those with a score above -3 are regarded as harmless [90]. PrimateAI is an algorithm based on deep learning that identifies harmful mutations in people with uncommon diseases [91]. This technique functions by removing prevalent missense variations found in six non-human primate species, as these mutations are predominantly clinically benign in humans [91]. MPC (missense badness, Polyphen-2, and constraint) is a logistic regression-based algorithm that identifies areas inside genes that are deficient in missense variants in ExAC data and utilizes variant-level metrics to estimate the effect of missense variations [92, 93]. A higher MPC score signifies more deleteriousness of amino acid modification in missense-constrained locations [92]. GenoCanyon is a whole-genome anno-tation-based tool that employs unsupervised statistics leveraging 26 computational and experimental annotations to predict the deleteriousness of mutations and the functional potential of each genomic position [94].

### 2.2 Molecular dynamics (MD) simulation

The crystal structure of the IAPP with bearing ID of 2L86 was retrieved from the RCSB protein databank, and this structure was then prepared by harnessing Schrödinger 2023-2 (Schrödinger, LLC, New York, NY, USA) according to the previously described protocols [95-100]. Following the preparation, we utilized the same software to construct the variants (T37P, L45P, G66R, T69I) harnessing residue scanning calculation program in Schrödinger 2023-2 [95, 101, 102]. Besides, we also generate the clinically found mutant S53G (same as S20G) through the residue scanning calculation program in Schrödinger 2023-2 to compare with these variants and the wild type [95, 101, 102].

MD simulations were executed utilizing the academic version of the Desmond program Schrödinger software 2023-2 (Schrödinger, LLC, New York, NY, USA) [103, 104] with the aim to apprehend the conformational dynamics of IAPP impaired as a consequence of variation. The OPLS4 force field was implemented to model the structures. The structures were placed in an orthorhombic box with dimensions of 10 Å in each direction [97, 105-108]. The explicit solvation model was applied; specifically, the Monte Carlo simulated transferable intermolecular potential 3 points (TIP3P) water model. In order to neutralize the systems, additional counterions (Na^+^/Cl^-^) were introduced, and the salt content of the system was modified to 0.15 M to uphold physiological environments. Before executing the MD simulations, each solvated system experienced minimization and equilibration operations using the default Desmond protocol. This protocol includes a sequence of controlled minimization steps and MD simulations, according to the report discussed earlier [109]. The simulation was executed using thermodynamic conditions, and to maintain a pressure of 1 atm and a temperature of 300 K, the isotropic Martyna-Tobias-Klein barostat [110] and the Nose-Hoover thermostat [111] were employed. The particle mesh Ewald technique was used to determine the long-range electrostatic interactions, whereas a cut-off of 9.0 Å was used to assess the short-range electrostatic interactions. The multistep RESPA integrator was employed to combine the equations of motion for both bound and non-bonded interactions within a specified short-range cut-off [112]. The integration was performed using an inner time step of 2.0 fs. A time step of 6.0 femtoseconds was used for non-bonded interactions that extended beyond the cut-off distance. Lastly, employing the NPT ensemble method, each simulation was conducted for a period of 200 ns 5 times for each system, and after each 100 ps, the coordinates were recorded. Following this, the stability of each system was assessed using MD simulation trajectories, employing RMSD (Root Mean Square Deviation), Rg (Radius of Gyration), and SASA (Solvent Accessible Surface Areas), RMSF (Root Mean Square Fluctuation), through Schrödinger 2023-2.

## 3. RESULTS

### 3.1 Screening and identification of deleterious SNPs

Accumulating studies have suggested that mutation in the *IAPP*, notably in the amyloidogenic segment (20-29 amino acid residues), causes its aggregation and amyloid formation, which leads to β-cell toxicity and death in type 2 diabetes mellitus (T2DM) [42-44, 64] and protein misfolding disorders (PMDs) [65, 66]. Therefore, a total of 62 missense variants associated with *IAPP* gene were extracted from the NCBI dbSNP database [67] and subsequently subjected to prediction for deleteriousness using 26 distinct *in silico* tools, including SIFT, FATHMM, M-CAP, MetaLR, MutPred, MutationTaster, PROVEAN, PolyPhen-2 HumDiv and HumVar, VEST4, fathmm-MKL, Condel, PhD-SNP, SNPs&GO, I-Mutant 3.0, CADD-PHRED and CADD-Raw, ClinPred, DANN, MVP, MetaRNN, REVEL, PredictSNP, LIST-S2, PANTHER, PrimateAI, MPC and GenoCanyon. These tools predict the deleteriousness of variations based on either sequence or a combination of sequence and structure (**Table S1, Supplementary File 2**). The total number of nsSNPs was 62, as shown in **Figure 2B**, where MutationTaster, CADD (PHRED, Raw), DANN, MPC, PredictSNP, and MetaRNN were able to detect most of them as deleterious, with MutationTaster capturing 51 nsSNPs and DANN and CADD-PHRED capturing 50 nsSNPs each. CADD-Raw and MPC could detect 48 nsSNPs, while PredictSNP and MetaRNN captured 46 and 45, respectively, as deleterious. Conversely, MetaLR, MutPred, GenoCanyon VEST4, and PolyPhen-2 HumVar predicted the lowest number of nsSNPs, while FATHMM, PANTHER, and PrimateAI captured none. The correlation between the used algorithms was represented in **Figure 2C**, where each tool varied the significance value. Most tools displayed an almost positive correlation except for FATHMM, PANTHER, and PrimateAI, which showed no correlation. Predicted SNPs, deemed deleterious by the most used algorithms, are usually more likely to be deleterious [113]. Therefore, nsSNPs that were assessed deleterious by at least eighteen algorithms were considered high-risk nsSNPs, such as rs1484149172 (L45P), rs374152437 (L6Q), rs78822118 (T37P), rs200376097 (G66R), rs758292552(L20P), rs763604023 (T69I). However, the variants L6Q and L20P were outside the mature IAPP protein sequence (34 – 70), and thus, the other four variants (T37P, L45P, G66R, T69I) were considered deleterious and selected for further study by molecular dynamics (MD) simulation (**Table S2, Supplementary File 2**).

**Figure 1.**
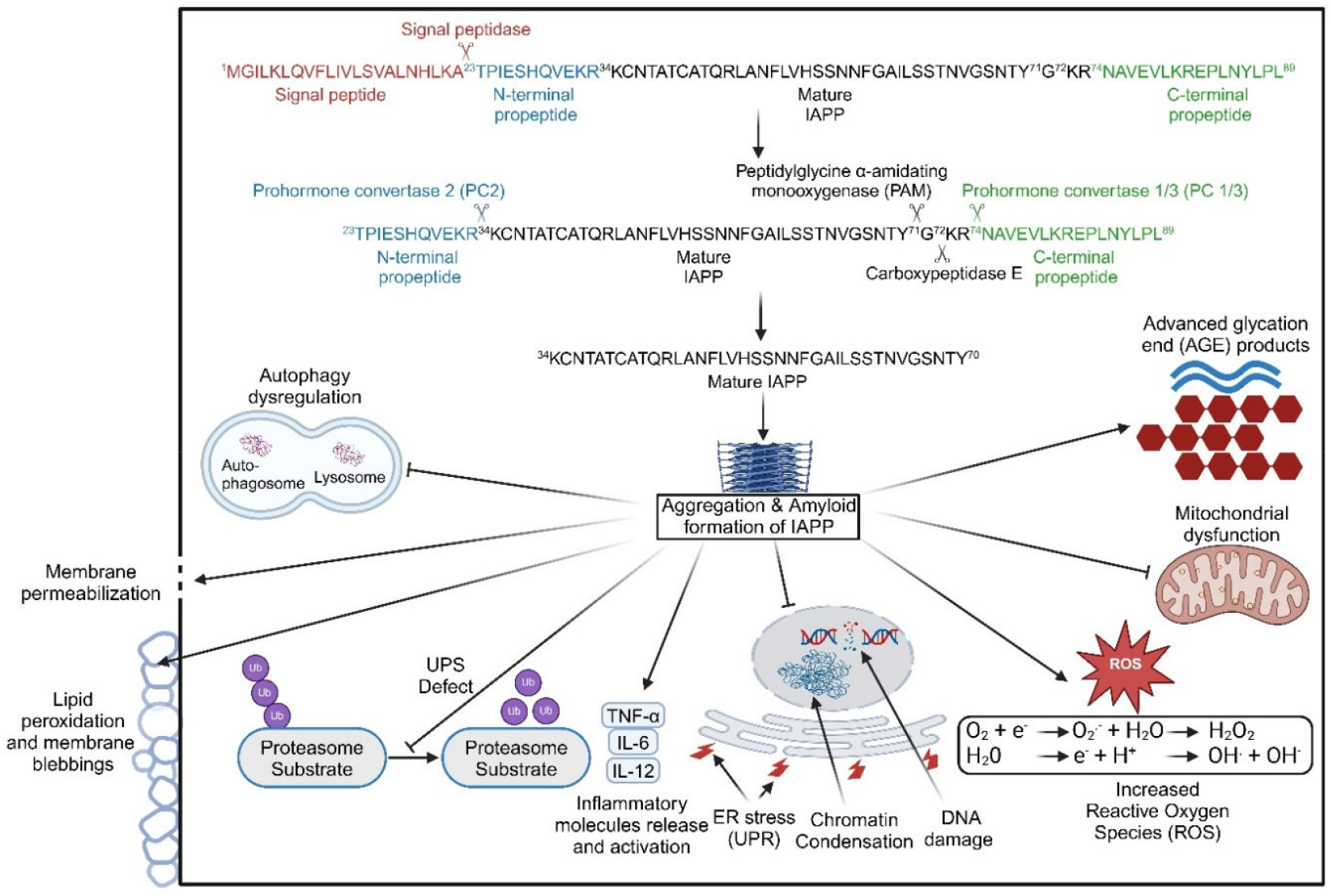
Processing of IAPP following translation by ribosome. *IAPP* gene is translated into pre-pro-IAPP (89 amino acid residues), which is then converted to 69 residues pro-IAPP by signal peptidase, and pro-IAPP is in turn processed to mature IAPP by PC2, PC1/3, CPE, and PAM. IAPP, in its aggregated and amyloid forms, induces cytotoxicity and β-cell death through mechanisms such as membrane permeabilization, plasma membrane blebbing, chromatin condensation, DNA fragmentation, elevated endoplasmic reticulum and oxidative stress, proinflammatory cytokine release and activation, increased reactive oxygen species (ROS) production, mitochondrial dysfunction, and the formation of advanced glycation end products. IAPP is also responsible for autophagy dysregulation and UPS defects.

**Figure 2.**
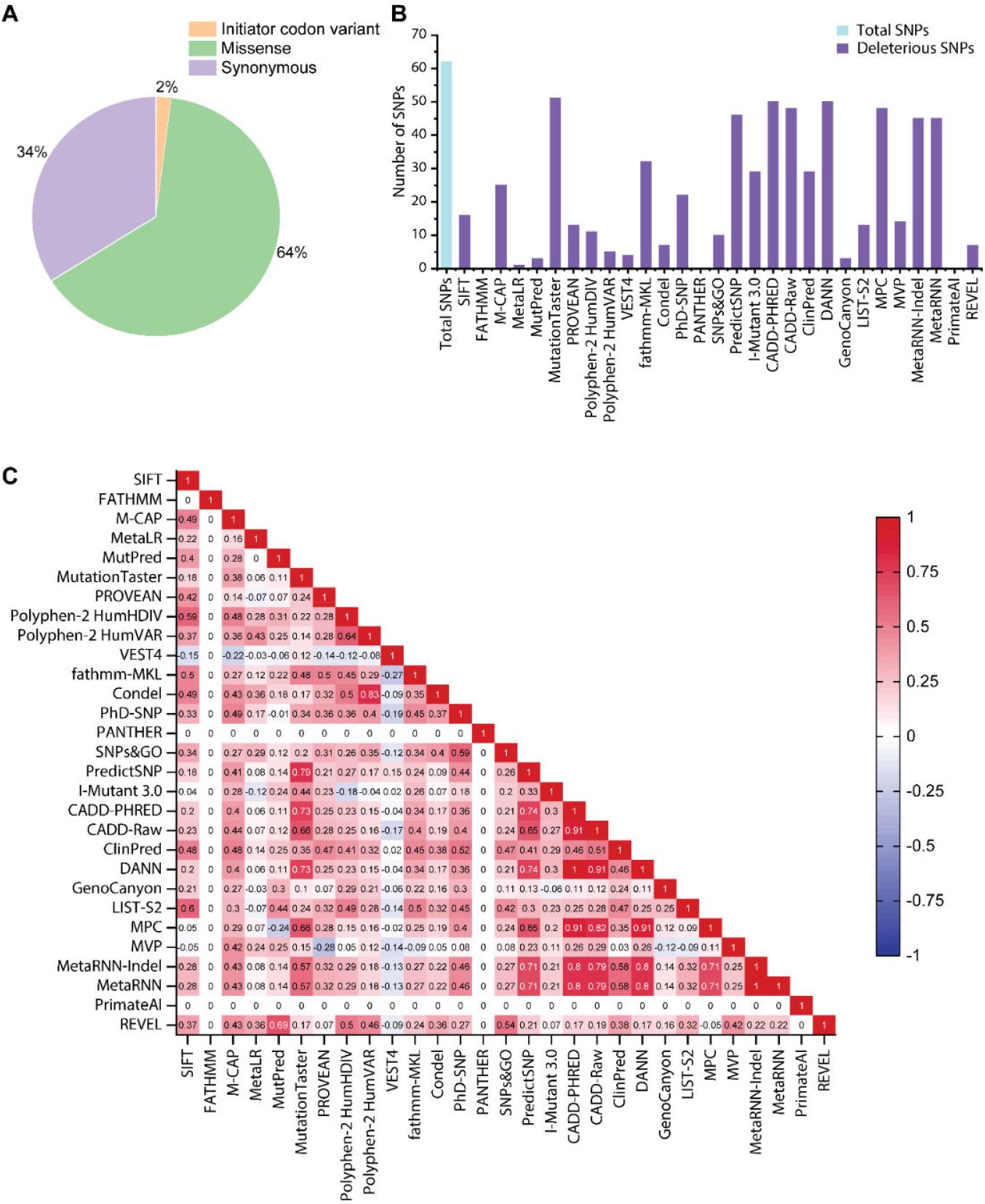
Extraction and prediction of deleterious non-synonymous SNPs in the *IAPP* gene. (A) The pie chart shows the percentage of different SNPs, with missense being 64% of total SNPs. The number of initiator codon variant, missense, and synonymous SNPs were 2, 78, and 41, respectively. Among the 78 missense SNPs, 62 SNPs were unique. (B) The bar plot demonstrates the total number of nsSNPs predicted by distinct computational tools. (C) The pairwise correlation for deleterious nsSNPs prediction among the tools is shown by a colorcoded heatmap (where blue, white, and red indicate negative, neutral, and positive correlation, respectively).

### 3.2 Molecular dynamics (MD) simulation

To elucidate the structural dynamics of IAPP resulting from variants, MD simulation on a microsecond timescale studied four predicted deleterious variants (T37P, L45P, G66R, and T69I) and the wild IAPP. In addition to these structures, S53G was also considered a negative control as it causes aggregation and amyloid formation of IAPP. Each structure underwent five separate MD runs of 200 ns each (1 μs in total). Afterward, the root mean square deviation (RMSD) analysis of each run for all structures was plotted, as shown in **Figure S1 (Supplementary File 2)**, to evaluate the equilibrated trajectories by considering the initial protein backbone structures. The results displayed that all structures were stable throughout the simulation. Thus, the 200 ns trajectory data from each run was considered and concatenated to generate 1 μs sub-trajectory for further analysis to maintain conformation sampling efficiency in the simulated trajectory analysis.

Next, the RMSD, Rg, SASA, and RMSF plotted were generated utilizing 1 μs trajectory data as shown in **Figure 3 (A-D)**. L45P and G66R significantly augmented the RMSD value compared to the wild type and even S53G, disrupting the flexible nature of IAPP and promoting dynamic behaviors, while T37P reduced the RMSD value, reducing the IAPP dynamic behavior. Unlike other variants, T69I could not notably alter this value, indicating that it did not impair IAPP’s flexible and dynamic nature (**Figure 3A**). In contrast, only G66R notably reduced the compactness of IAPP, as evidenced by its extensive Rg value, indicating that it enhanced the dynamic behavior of IAPP, compared to other variations (L45P, T69I) and control (S53G). T37P lowered the Rg value, but this change was not significant. The Rg value of wild was 10.58 Å and ranged from 8.83 Å to 19.68 Å, consistent with the previous study where the authors predicted Rg value ranging from 8.3 to 13.9 Å [114]. The same sequence was also observed in the case of SASA, where only G66R elevated the SASA value compared to other variants and control (S53G), suggesting that G66R significantly augmented the hydrophobic surface area of IAPP accessible to the solvent, thus disrupting the stability of IAPP. Overall, RMSD, Rg, and SASA data indicated that G66R and L45P may affect the conformational deviations of IAPP; additionally, T37P may affect them to some extent.

**Figure 3.**
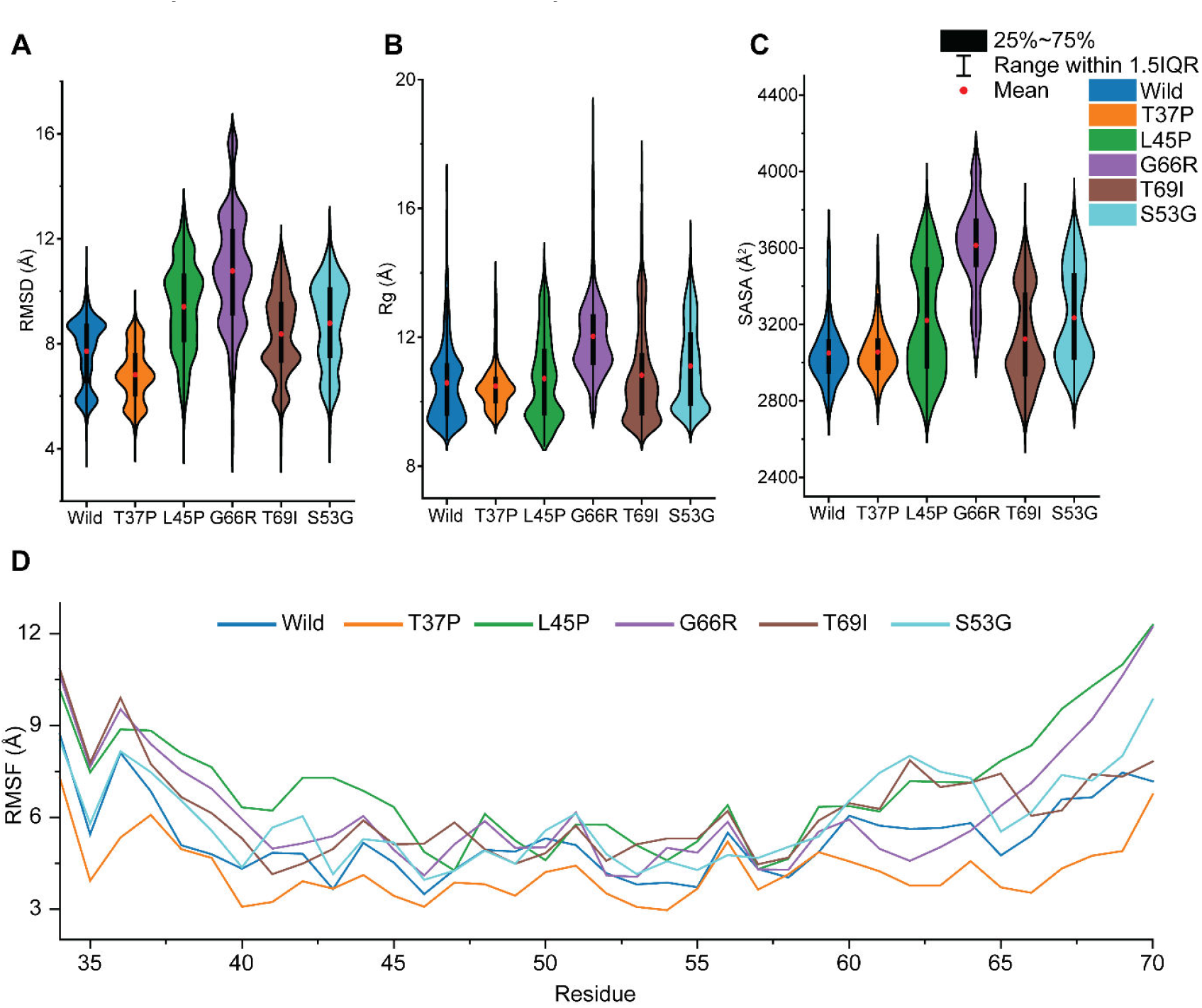
Variants alter the conformational dynamics of IAPP. The graphical representation in violin plots depicts the mean difference in conformational alterations between wild-type, variant-containing structures (T37P, L45P, G66R, and T69I) and negative control (S53G) using RMSD (A), Rg (B), and SASA (C) assessment. The red circle, error bars, and black dots in the violin plots (A-C) indicate the mean, standard deviation (SD), and individual RMSD (A), Rg (B), and SASA (C) values, respectively. (D) The line plot represents the variations in residual flexibilities among wild variants (T37P, L45P, G66R, and T69I) and negative control (S53G), plotted by the Cα-root mean square fluctuation (RMSF).

Then, we assessed the effect of variation on the local fluctuation of IAPP by considering individual amino acid fluctuation. As shown in **Figure 3D**, all variants altered IAPP backbone fluctuation as indicated by changes in the amino acid positions. L45P induced more amino acid fluctuations than other variants and wild types across the local amino acids. Conversely, T37P lowered the fluctuation pattern of amino acids, consistent with RMSD, Rg, and SASA. However, unlike RMSD, Rg, and SASA data, G66R showed less deviation in amino acid fluctuation. All variants altered amino acid fluctuation patterns in the disulfide bond region 35-40 residues (2-7 residues), which is responsible for maintaining native α-helical structure [115]. This fluctuation may reduce helical structure and promote β-sheet content, forming IAPP amyloid fibril [115]. Additionally, the amyloidogenic region 53-62 (20-29 residues) were altered in fluctuation due to all variants, thus may induce IAPP to adopt the β-sheet structure found in amyloid fibril formation, consistent with the previous study [116, 117]. It has been reported that Gln43, Leu45, Asn47, and Leu49 form contact with each other and with Arg44, Ala46, and Phe48, thus stabilizing the β-sheet structure formation during amyloid fibril formation [118]. Our study observed that amino acids fluctuated less within these residues, notably within Ala46, Asn47, Phe48, and Leu49. Overall, RMSF data indicate that all variants alter the conformational dynamics of IAPP, most notably T37P, L45P, and G66R.

## 4. DISCUSSION

Mutation in IAPP, especially in the amyloidogenic region has been reported to cause aggregation and amyloid formation, which causes cytotoxicity, β-cell death (apoptosis) [42-44, 64], and protein misfolding disorders and neurodegenerative disorders, which includes, Alzheimer’s disease (AD), Parkinson’s disease (PD), amyotrophic lateral sclerosis (ALS), and prion disorders [65, 66]. The current was intended to unveil the deleterious variations in *IAPP* that might cause aggregation and amyloid formation. Here, we revealed three deleterious nsSNPs (T37P, L45P, and G66R) that conformation dynamics of IAPP may result in IAPP aggregation and amyloid formation, but a detailed study is needed.

The current study first performed predictions for deleterious variations of *IAPP* using the 62 nsSNPs found by the NCBI dbSNP database [67]. We harnessed 26 widely used *in silico* tools for deleteriousness prediction. Subsequently, the top 4 variants, predicted deleterious by at least 18 tools, were deemed the most harmful variants and, thus, considered for further assessment by MD simulation on a microsecond scale. The RMSD, Rg SASA, and RMSF data indicated that T37P, L45P, and G66R were the most deleterious variants which affected the conformational alterations of IAPP. The RMSF data suggested that all variants showed notable fluctuation with the disulfide bond region, responsible for maintaining α-helical contents [115] and within the amyloidogenic region, accounting for amyloid formation [116, 117]. Therefore, variation in *IAPP* (notably T37P, L45P, and G66R) may cause its aggregation and amyloid formation; however, a more detailed study is indispensable to probe this.

## 5. CONCLUSION

Understanding the consequences of variants on the conformational dynamics of IAPP is essential, as emerging studies indicate that variations in *IAPP*, particularly within the amyloidogenic region, lead to its aggregation and amyloid formation, resulting in β-cell toxicity and mortality in type 2 diabetes mellitus (T2DM) and protein misfolding disorders (PMDs). Considering this, the current work, by employing extensive bioinformatics and MD simulation studies, identified 3 variants in *IAPP* (T37P, L45P, and G66R) that alter the conformational dynamics of IAPP, which might be associated with IAPP aggregation and amyloid formation. Therefore, this study can be a starting point for assessing the consequences of the identified variants on IAPP aggregation and amyloid formation by in vitro and/or in vivo studies and designing novel therapeutics as potential treatments against IAPP aggregation and amyloid formation, hence preventing T2DM and PMDs.

