## Supplementary material for "Elucidating conformational alteration of human islet amyloid polypeptide by nonsynonymous substitution": (Supplementary File 2)

**Table S1.** The cut-off of predicted computational tools employed for identifying harmful nonsynonymous single nucleotide polymorphisms (nsSNPs) in the *IAPP* gene.

| SL.NO | Tool name | Prediction method | Cut-off value |
| --- | --- | --- | --- |
| 1 | SIFT | Sequence | ≤0.05 |
| 2 | FATHMM | Sequence | >0.5 |
| 3 | M-CAP | Sequence | >0.025 |
| 4 | MetaLR | Sequence | >0.5 |
| 5 | MutPred | Sequence and structure | >0.75 |
| 6 | MutationTaster | Sequence | >0.5 |
| 7 | PROVEAN | Sequence | ≤-2.5 |
| 8 | Polyphen-2 | Sequence and structure | ˃0.9 |
| 9 | VEST4 | Sequence | <0.05 |
| 10 | fathmm-MKL | Sequence | >0.5 |
| 11 | Condel | Sequence | >0.9 |
| 12 | PhD-SNP | Sequence | >0.5 |
| 13 | PANTHER | Sequence | <-3 |
| 14 | SNPs&GO | Sequence | >0.5 |
| 15 | PredictSNP | Sequence and structure | <1 |
| 16 | I-mutant 3.0 | Sequence | <-0.5 |
| 17 | CADD | Sequence | >0.5 |
| 18 | ClinPred | Sequence and structure | >0.05 |
| 19 | DANN | Sequence | >0.5 |
| 20 | GenoCanyon | Sequence | >0.1 |
| 21 | LIST-S2 | Sequence | >0.85 |
| 22 | MPC | Sequence | <1 |
| 23 | MVP | Sequence and structure | >0.75 |
| 24 | MetaRNN | Sequence and structure | >0.05 |
| 25 | PrimateAI | Sequence | >0.8 |
| 26 | REVEL | Sequence | Between 0.5 and 0.75 |

**Table S2.** Summary of top 4 deleterious nsSNPs predicted by different computational tools.

| rs ID | rs1484149172 | rs200376097 | rs763604023 | rs763676184 | rs78822118 |
| --- | --- | --- | --- | --- | --- |
| Substitution | L45P | G66R | T69I | R44S | T37P |
| Tools predicted deleterious | 23 | 18 | 18 | 17 | 19 |
| SIFT | 0 | 0 | 0 | 0 | 0 |
| FATHMM | 0.88 | 1.56 | 1.74 | 1.85 | 1.71 |
| M-CAP | 0.039 | 0.159 | 0.049 | 0.031 | 0.069 |
| MetaLR | 0.127 | 0.246 | 0.151 | 0.064 | 0.132 |
| MutPred | 0.85 | - | 0.518 | 0.712 | 0.735 |
| MutationTaster | 1 | 1 | 1.000 | 0.998 | 1.000 |
| PROVEAN | -6.86 | -7.07 | -3.04 | -5.47 | -5.55 |
| Polyphen-2 HumDIV | 1 | 0.96 | 0.997 | 0.088 | 0.999 |
| Polyphen-2 HumVAR | 1 | 0.808 | 0.955 | 0.186 | 0.974 |
| VEST4 | 0.984 | 0.952 | 0.68 | 0.717 | 0.603 |
| fathmm-MKL | 0.966 | 0.972 | 0.941 | 0.678 | 0.928 |
| Condel | 0.945 | 0.748 | 0.843 | 0.462 | 0.861 |
| PhD-SNP | 0.768 | 0.731 | 0.648 | 0.773 | 0.648 |
| PANTHER | 0.776 | 0.732 | 0.479 | 0.39 | 0.518 |
| SNPs&GO | 0.707 | 0.695 | 0.461 | 0.618 | 0.346 |
| PredictSNP | 0.869 | 0.869 | 0.756 | 0.869 | 0.756 |
| I-Mutant 3.0 | -0.65 | - | 0.67 | -0.64 | -1.47 |
| CADD-PHRED | 27.1 | 27.1 | 23.6 | 22.1 | 24.6 |
| CADD-Raw | 4.011 | 4.015 | 3.106 | 2.398 | 3.462 |
| ClinPred | 0.998 | 0.996 | 0.924 | 0.992 | 0.996 |
| DANN | 0.999 | 0.999 | 0.997 | 0.965 | 0.988 |
| GenoCanyon | 0.826 | 0.044 | 0.042 | 2.355 | 3.635 |
| LIST-S2 | 0.893 | 0.941 | 0.917 | 0.934 | 0.877 |
| MPC | 0.653 | 0.562 | 0.366 | 0.489 | 0.585 |
| MVP | 0.724 | 0.553 | 0.586 | 0.346 | 0.587 |
| MetaRNN-Indel | 0.957 | 0.796 | 0.586 | 0.666 | 0.836 |
| MetaRNN | 0.963 | 0.802 | 0.450 | 0.594 | 0.845 |
| PrimateAI | 0.555 | 0.595 | 0.442 | 0.264 | 0.358 |
| REVEL | 0.689 | 0.65 | 0.216 | 0.258 | 0.449 |

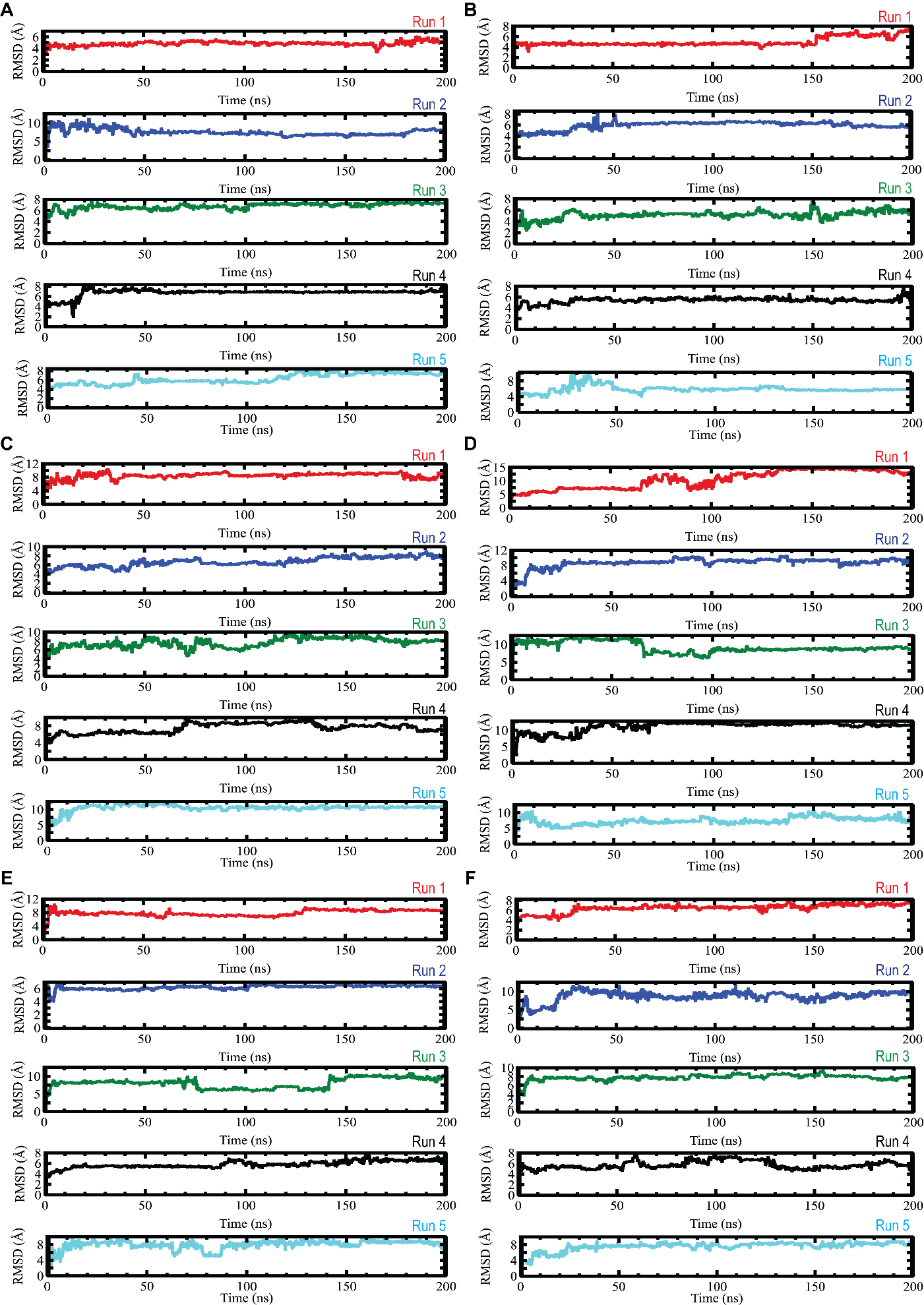

**Figure S1.** Line plots showing RMSD values produced from each run of the particular structure (wild, variants and negative control), were assessed utilizing C-alpha of wild (A), T37P (B), L45P (C), G66R (D), T69I (E), and negative control S53G (F), respectively.
